# Poly(ADP-ribose) binding sites on collagen I fibrils for nucleating intrafibrillar bone mineral

**DOI:** 10.1101/2024.07.02.600619

**Authors:** Marco A. Zecca, Heather F. Greer, Melinda J. Duer

## Abstract

Bone calcification is essential for vertebrate life. The mechanism by which mineral ions are transported into collagen fibrils to induce intrafibrillar mineral formation requires a calcium binding biopolymer that also has highly selective binding to the collagen fibril hole zones where intrafibrillar calcification begins, over other bone extracellular matrix components. Poly(ADP-ribose) has been shown to be a candidate biopolymer for this process and we show here that poly(ADP-ribose) has high affinity, highly conserved binding sites in the collagen type I C-terminal telopeptides. The discovery of these poly(ADP-ribose)-collagen binding sites gives new insights into the chemical mechanisms underlying bone calcification and possible mechanisms behind pathologies where there is dysfunctional bone calcification.

## Introduction

The deposition of solid calcium phosphate-based mineral within bone extracellular matrix is an essential physiological process. The mineral forms inside and between collagen fibrils (1), but how collagen fibrils are selected for calcification over other nanostructures or proteins in the extracellular matrix (ECM) is still an open question. It has been extensively hypothesized that the selection is via one or more biomolecules that bind calcium ions and collagen, thus delivering the calcium ions to the correct location for mineral formation. This hypothesis was later refined by postulation of the polymer-induced liquid precursor model (2), whereby a polyanionic biomolecule not only delivers calcium ions specifically to collagen fibrils, but establishes a calcium-rich, fluid, amorphous precursor mineral phase which flows into collagen fibril hole zones, the fibril region where mineral nucleation has been evidenced by transmission electron microscopy (TEM) to begin (3–6). This latter model can account for mineral forming inside collagen fibrils, as well as between them, and for the close fitting of mineral crystals to the spaces available inside the fibrils (7). What is missing from this mechanism of collagen fibril calcification is the identity of the biomolecule responsible for the delivery of calcium ions to collagen fibril hole zones. The biomolecule must bind calcium ions in abundance to deliver sufficient for mineral nucleation at least, must bind to the collagen fibril hole zone where mineral formation begins and likely establishes a fluid amorphous mineral precursor phase suggesting that the biomolecule is polyanionic. Several extracellular proteins expressed by osteoblasts, the bone-forming cells, have been extensively studied but none have acquired sufficient evidence to be considered as plausible candidates (8– 10).

Previous work has shown that the extracellular concentration of a polyanionic nucleotide, poly(ADP ribose) (PAR) correlates with both bone and vascular calcification (11), and that inhibition of PAR production inhibits calcification by osteoblasts and vascular smooth muscle cells (VSMCs) *in vitro* (12). Poly(ADP-ribose) (PAR) is a post-translational modification added to proteins by PAR polymerases (PARPs). Expression of PARP1, the main PAR-producing enzyme, has been shown to be essential for osteoblast matrix calcification in vitro (13) and important in pathological vascular calcification (12, 14)

Importantly, PAR has been observed to form dense liquid droplets with calcium *in vitro*, with high selectivity for calcium over other divalent metal cations and so may be capable of generating a fluid mineral precursor phase in bone calcification in vivo. PAR-Ca droplets were also shown to bind to collagen I fibrils (12), with selective affinity for the fibril hole zones, further marking PAR out as a potential biomolecule for initiating collagen calcification in bone.

Here, we explore the hypothesis that PAR-Ca droplets initiate intrafibrillar mineralization by PAR-Ca droplets by having high affinity binding sites in collagen I hole zones and elucidate the PAR binding site on collagen type I fibrils.

## Results

### TEM shows PAR-Ca droplets selectively bind to the collagen fibril *a* sub-band

We began by examining in more detail where PAR-Ca droplets bind on collagen fibrils. A suspension of PAR-Ca droplets was added to collagen fibrils on a TEM grid and allowed to remain undisturbed for 60s, then uranyl acetate stain was added and the system left undisturbed for a further 40s, after which the suspension was blotted off the grid. In TEM images of these samples, PAR-Ca droplets were seen bound to collagen fibrils, with significant preference for the *a* sub-band (Fig. 1A,B,E, Fig S1A,B). Most cases of binding to other sub-bands were seen when nearby *a* sub-bands were already occupied by PAR-Ca droplets. Almost no PAR-Ca droplets were seen binding at overlap zones (the *b* sub-band), and very few unbound PAR-Ca droplets were found on the grids, indicating that the PAR-Ca droplets have a high binding affinity for collagen fibrils.

**Figure 1:**
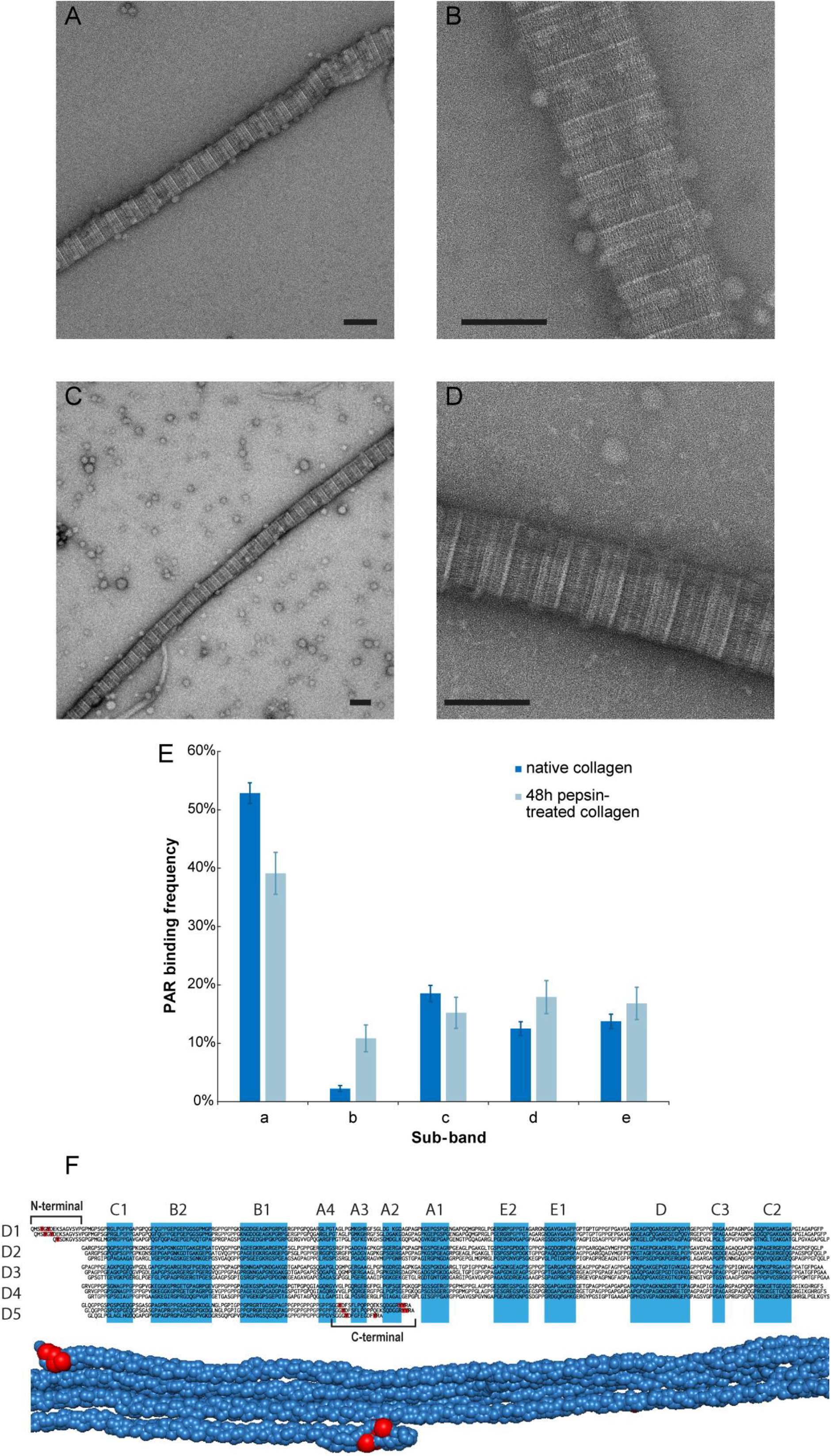
A-D: Representative TEM images of PAR-Ca droplets bound to an untreated collagen fibril (A,B) and a collagen fibril formed from collagen that had been degraded with pepsin for 48 hours prior to fibrillization (C,D), showing decreased binding affinity of PAR-Ca droplets to pepsin-digested collagen fibrils (scale bars 100 nm); E: Proportion of PAR-Ca droplets bound at each collagen sub-band in TEM images of PAR-Ca droplets and raw and pepsin degraded collagen fibrils, p<0.001; F: Schematic of the amino acid sequence of a collagen I fibril (Mus musculus) with sub bands and tyrosine residues highlighted.

The collagen I fibril *a* sub-band where the PAR-Ca droplets preferentially bind contains the C-terminal telopeptides. To test whether the C-terminal telopeptides are involved in PAR-Ca droplet binding, pepsin was used to remove both the C- and N-terminal telopeptides from collagen molecules prior to fibril formation.

Fibril formation was found to be significantly slower after incubation with pepsin, consistent with telopeptide removal (15): 72 hours at 4°C for visible fibrils to form after 48 hours of pepsin treatment compared to 12-24 hours for untreated collagen.

Removal of telopeptides led to a significant decrease in PAR-Ca droplets binding to collagen fibrils in TEM images, with many PAR-Ca droplets now found on the grids unbound to collagen fibrils (Fig. 1C,D, Fig. S1C,D). In the limited droplet-fibril binding that was observed, there was statistically significant (p<0.001) reduced selectivity for the *a* sub-band compared to normal collagen fibrils (Fig. 1E). We note that pepsin digestion is unlikely to result in complete removal of all telopeptides and therefore that some of the residual binding observed could potentially be attributed to fibrils with only partial telopeptide removal. Taken together, the data suggest that the primary PAR-Ca droplet binding sites are situated in the C-terminal telopeptides of collagen I, or otherwise their presence is required to induce binding.

### The collagen type I C-terminal telopeptides contain highly conserved PAR binding sequences

Several consensus PAR binding motifs on proteins are known in biology (16). We analysed the vertebrate consensus amino acid sequences of both the α1 and α2 chains of collagen I for regions with similarity to any of these known PAR binding motifs. The only sequences in either chain with similarity to any of the known PAR binding sequences are in the C-terminal telopeptides, where we found in both the α1 and α2 chains, Y…YR motifs similar to the consensus PBZ PAR binding motif (17, 18) (Fig. 1D).

Conservation analysis over vertebrate collagen type I sequences revealed that the C-terminal telopeptides of both α1 and α2 chains of collagen I are very highly conserved (Fig. 2A,B), particularly if the substitution of tyrosine for phenylalanine and glutamate for aspartate are considered as contributing similar chemical functionality. Both C-terminal telopeptides are more than 90% conserved at most residues, suggesting that they must contain regions of significant biological importance.

**Figure 2:**
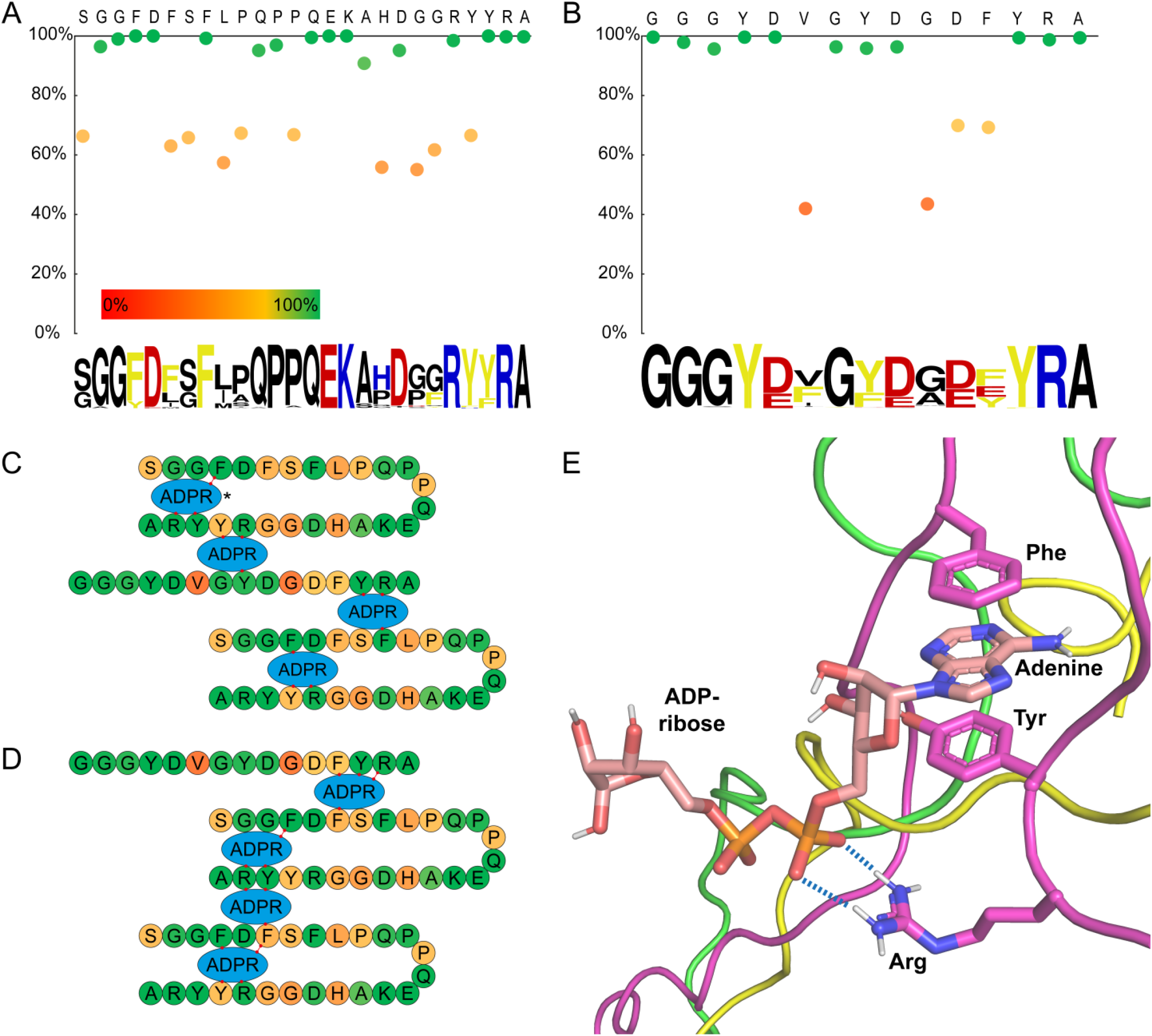
A, B: Conservation plots and sequence logos of the C-terminal telopeptides of α1 (including α1a for Actinopterygii) (A) and α2 (B). Percentage conservation was calculated accounting for equivalence between the chemically functionally similar aspartate and glutamate, and between phenylalanine and tyrosine; C, D: Schematic representations of potential binding sites for ADP-ribose units between chains in the tropocollagen C-terminal, coloured according to conservation at each site; E: potential binding mode of a single ADP-ribose molecule to a C-terminal telopeptide model based on the X-ray structure determined by Orgel et al. (19), the binding mode shown is indicated by *.

In the known PBZ motifs, the arginine and tyrosine residues are essential for PAR binding, and these residues are particularly highly conserved in the C-terminal telopeptides of both collagen type I α1 and α2 chains across all vertebrates.

The binding mode of PAR in the PBZ motif in aprataxin PNK-like factor (APLF) has been shown to be *via* π-stacking interactions between the aromatic rings of two conserved tyrosine residues and the adenine moiety of an ADP-ribose unit, whilst a nearby arginine side chain engages in a charge-charge interaction with the pyrophosphate group of the neighbouring ADP-ribose unit in the PAR polymer (18). The folding of the three chains of the collagen I heterotrimer suggested by the collagen I fibril X-ray diffraction structure (19), yields a number of similarly-organized possible PAR binding sites, both within single telopeptide a chains and between two telopeptide chains (fig. 2C-E). The presence of multiple potential binding sites in close proximity in other proteins has been shown to increase affinity for PAR, with the presence of two tandem PAR binding sites in APLF increasing its affinity for PAR over 500-fold relative to each of the two binding sites individually (20). This would suggest that the C-terminal telopeptide system may have a very high affinity for PAR within each tropocollagen molecule.

The regular, periodic nature of collagen molecules within collagen fibrils leads to regions spanning the circumference of the fibril in which numerous telopeptides are found in close proximity. From the X-ray diffraction structural model of collagen type I fibrils, the C-terminal telopeptides are expected to be separated by between ∼2.7 and 4.0 nm (the *a* and *b* dimensions of the collagen fibril crystallographic unit cell (19)) and the length of a PAR monomer unit is of order 1.5 nm (from molecular modelling). Thus, a 100-monomer unit PAR polymer could potentially contact up to 50 collagen binding sites. This multitude of potential PAR binding sites on a collagen fibril could be expected to significantly increase the affinity of PAR to collagen fibrils compared to isolated collagen molecules and account for the high affinity of PAR-Ca binding to collagen fibrils we observed by TEM.

### Solution-State NMR shows C-terminal telopeptide Tyr residues π-stack with ADP ribose adenine rings

Having identified possible binding sites in the C-terminal telopeptides of both the α1 and α2, we tested each of the telopeptides for their respective affinities to bind PAR. Short, water-soluble peptides comprising the human sequences of each of the C-terminal telopeptides were used to investigate binding by solution-state NMR, a powerful method to simultaneously identify peptide binding sites and quantify their binding equilibria in aqueous conditions. The lengths and extents of branching of PAR polymers involved in bone calcification *in vivo* are not known, so the simplest possible model for PAR was used: monomeric ADP-ribose. Our rationale was that if the ADP-ribose monomer can be shown to bind to a single C-terminal telopeptide a chain, even weakly, then a PAR polymer with many ADP-ribose subunits can be expected to bind more strongly to the C-terminal telopeptides of a full heterotrimeric tropocollagen where there will additionally be multiple ADP-ribose binding sites. To identify which residues of the model peptides are involved in any binding to ADP-ribose, we followed the changes in ^1^H chemical shifts for both the peptides and ADP-ribose upon titration of ADP-ribose into solutions of each peptide, at three different temperatures. ^1^H chemical shifts are sensitive to changes in peptide molecular conformation induced by ligand binding to them; specifically, the π-stacking binding modes hypothesized to take place between the C-terminal telopeptides and ADP-ribose are expected to result in significant changes in the Tyr aromatic ring ^1^H chemical shifts, and similarly for any arginine residues involved in ADP-ribose binding. Varying the temperature of the peptide-ADP-ribose solution is expected to shift any binding equilibrium which should be manifest by a clear temperature dependence of the ^1^H chemical shift changes.

A combination of COSY, TOCSY, NOESY and ROESY NMR spectra were used to assign the proton signals for the two model peptides to the respective amino acid residues in each peptide. In particular, ^1^H signals corresponding to the tyrosine residues at sites 23 and 24 were distinguished for the α1 model peptide, and those corresponding to the tyrosine residues at sites 7, 11, and 16 were distinguished for the α2 model peptide (Table S1).

Upon addition of ADP-ribose to the buffered peptide solutions, the ^1^H chemical shift of all tyrosine aromatic protons decreased (Fig. 3 A-H), consistent with a π-stacking interaction between the adenine ring of ADP-ribose and the aromatic rings of the tyrosine residues. The ^1^H chemical shifts of arginine δ protons decreased in the α1 model peptides by a similar magnitude to the tyrosine aromatic protons in the same peptide with increasing ADP-ribose concentration, whereas the ^1^H chemical shift of the arginine δ protons in the α2 model peptides decreased with increasing ADP-ribose concentration only at lower temperatures (Fig. S2). There was negligible pH change over the course of the titrations. All other peptide ^1^H signal chemical shifts changed by a much smaller amount, confirming that the tyrosine and arginine chemical shift changes are from peptide-ADP-ribose interactions. The magnitude of changes in tyrosine aromatic proton ^1^H chemical shift increased as the temperature was decreased, indicating more molecules in the bound state at lower temperatures, consistent with expectation for peptide-ligand interactions. The largest peptide ^1^H chemical shift change with ADP-ribose concentration was for the aromatic ring of Tyr23 in the α1 model telopeptide (-GR**<u>Y</u>**YRA), suggesting that this residue has the strongest interaction with ADP-ribose. In the α2 model peptide, Tyr7 (-GGG**<u>Y</u>**D-) aromatic ^1^H chemical shifts decreased by a significantly greater amount with ADP-ribose concentration compared to the other two tyrosine residues, suggesting that this tyrosine in the α2 chain is the aromatic residue most strongly involved in ADP-ribose binding.

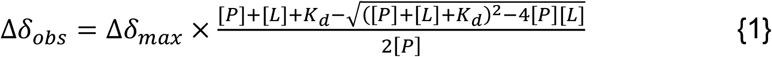

**Figure 3:**
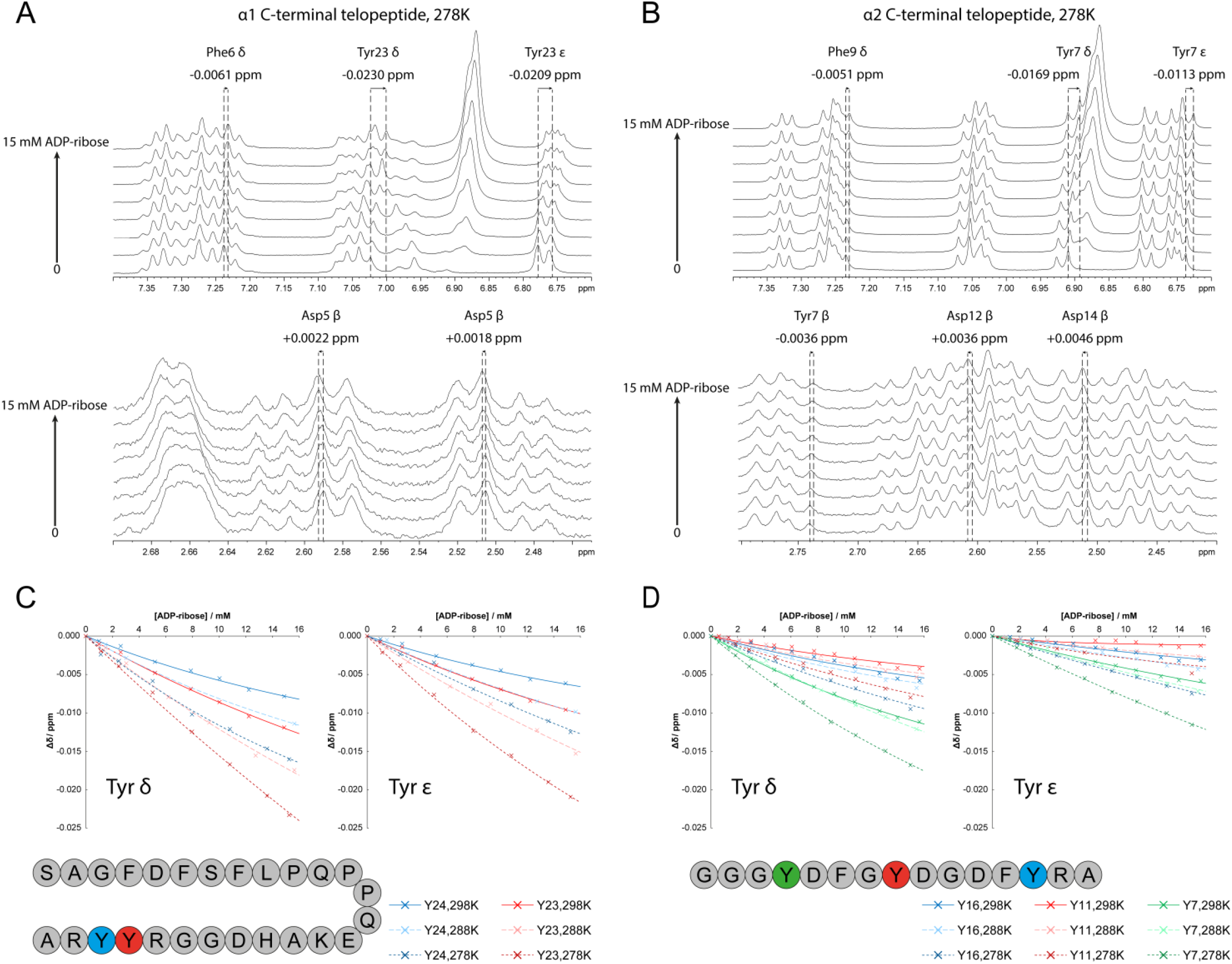
A,B: Sections of proton NMR spectra of the α1 (A) and α2 (B) at 278K showing changes upon addition of ADP-ribose. Representative chemical shift changes of individual peaks are shown over the full titration; C,D: Plots showing the decrease in chemical shift of tyrosine aromatic protons in the α1 (C) and α2 (D) model peptides upon addition of ADP-ribose. Approximate binding curves have been fitted according to equation {1}, with peptide and ligand ADP-ribose concentrations [P] and [L] in mM (21).

The adenine ^1^H chemical shifts of ADP-ribose are also sensitive to π-stacking, but this can include adenine rings π-stacking with each other. To account for this, the adenine ^1^H chemical shifts were plotted against the concentration of ADP-ribose for pure ADP-ribose and the data fitted to a straight line. The extrapolation of the line to zero ADP-ribose concentration gives the approximate adenine ^1^H chemical shift for an ADP-ribose molecule in the absence of any adenine-adenine π-stacking (Fig. S3 A,B). This adenine ^1^H chemical shift was compared to similar data extrapolations for the adenine ^1^H chemical shifts in the presence of 1 mM of each of the model peptides. The predicted ADP-ribose adenine ^1^H chemical shift in the absence of adenine-adenine π-stacking was lower in the presence of each of the model peptides, suggesting that in the presence of either telopeptide, the ADP-ribose adenine rings are involved in p-stacking interactions in addition to any adenine-adenine interactions. The predicted ADP-ribose adenine ^1^H chemical shift in the absence of adenine-adenine π-stacking was lower for the α1 model telopeptide, suggesting that adenine ring-peptide interactions are stronger for this telopeptide. At all temperatures, the decrease in adenine ^1^H chemical shifts caused by the presence of 1 mM of the α1 model peptide was approximately equivalent to that caused by 2 mM of adenine-adenine π-stacking, indicating particularly strong π-stacking between ADP-ribose and the α1 model peptide, suggesting high potential affinity of the α1 C-terminal telopeptide for PAR.

## Discussion

An important feature of collagen fibril calcification in bone is that mineral crystals are found both inside the fibril and extending between fibrils and this must be highly relevant to the mechanical properties of bone. Mineral ions must be deposited inside a collagen fibril before the extrafibrillar mineral forms, or the extrafibrillar mineral sheath will block the fibril to influx of the necessary ions for intrafibrillar mineralization. This implies that there must be a method of temporal control such that intrafibrillar mineralization is initially favored over extrafibrillar mineralization. One possible mechanism for this is the polymer-induced liquid-precursor (PILP) model for bone mineralization (2), which hypothesizes that a polyanion, with strong binding affinity for calcium ions causes a phase separation in the presence of Ca^2+^ ions, resulting in a calcium-rich, dense liquid precursor phase and that subsequent binding of the polyanion to collagen fibrils results in this precursor phase penetrating the collagen fibril hole zones and flowing into the spaces available for mineral crystal formation inside the collagen fibril (6). The authors of that seminal study (2) used polyaspartic acid as an exemplar polyanion; in vivo, such a process relies on a polyanion with strong, specific binding to collagen fibrils to ensure that it is collagen fibrils that are calcified and not any of the other bone extracellular matrix proteins. A key question in bone calcification then is: what is the bone matrix molecule that plays the role of polyaspartic acid *in vivo*?

Previous work has shown that PARP activity is required for osteogenic differentiation (22), and our previous work has shown that PAR has a strong, selective affinity for calcium ions and forms dense liquid droplets in the presence of Ca^2+^ concentrations below supersaturation (12). That same study showed biomimetic mineralization of collagen I fibrils in the presence of PAR-Ca droplets (12). Here we provide the final piece of evidence needed to demonstrate that PAR is a plausible candidate for the polyanion that in vivo forms precursor mineral and which initiates formation in intrafibrillar mineral crystals. We show here that PAR has multiple, closely-arranged binding sites in the C-terminal telopeptides of collagen type I, binding sites which are located precisely in the region of collagen fibrils where mineral nucleation has been shown to begin, namely the fibril hole zone. The amino acid sequences in the telopeptides that putatively bind PAR we show to be highly conserved across all vertebrates. Thus, these sequences must be involved in some essential processes or structures across a very wide range of animals. Bone calcification and its proper control are essential to life and occur across all vertebrates and could certainly be expected to drive high sequence conservation in protein regions governing mineral nucleation. That mineral nucleation begins *inside* collagen fibrils rather than on their surfaces calls for a robust chemical mechanism to ensure mineral ions are preferentially transported inside the fibrils rather than remaining in the vicinity of their surfaces, which the hydrophobic nature of collagen fibrils would tend to result in in the absence of a specific transport mechanism. Mutation of the final tyrosine residue in the α2 C-terminal telopeptide (in this work referred to as Y17) to cysteine has been found to correlate with cases of osteogenesis imperfecta (OI) (23), and several mutations and deletions of the key tyrosine and arginine residues in the α1 C-terminal telopeptide have also been proposed to correlate with OI (NIH ClinVar (24) accession codes: VCV000580848.5, VCV001486023.6, VCV001430512.4, VCV000456776.10, VCV001074867.6, VCV000930782.3, VCV000967962.6) further supporting the idea that these collagen C-terminal sequences are essential for functional bone calcification.

The periodic nature of collagen I fibrils with multiple, periodic PAR binding sites at the fibril hole zone suggests that the collagen fibril hole zone could bind PAR very strongly. This feature of periodic fibril sites binding a periodic polymer could drive selectivity over other potential PAR binding species in the ECM. Collagen fibrils are the only periodic structures in bone ECM and contain close to the maximum density of PAR binding sites that can possibly exist on any proteinaceous structure; even if another ECM protein has one or two strong PAR binding sites, it cannot match the strength of binding possible on a periodic collagen fibril. This concept gives rise to the intriguing question of how PAR-collagen binding and calcification might be affected if the collagen fibril structure is disrupted and whether such a mechanism could be at the root of some bone mineral pathologies.

The multiple binding sites throughout the interior of the collagen fibril hole zone may act to draw PAR-Ca liquid droplets into the fibril and assist their flow into the intrafibrillar spaces where mineral crystals will form. Strong binding at multiple sites throughout the fibril hole zone can expected to increase the free energy of the PAR-Ca liquid droplets compared to free PAR-Ca droplets in solution, through reduction of the PAR molecule conformational flexibility when it binds to collagen (reduction of entropy) and increase of the droplet interfacial energy as the droplet encounters the relatively hydrophobic collagen molecules (increase in enthalpy).

Increasing the free energy of the PAR-Ca droplet will in general lower the activation energy for mineral nucleation, suggesting that PAR-Ca droplet binding to collagen fibrils in the presence of inorganic phosphate could initiate mineral formation.

Several extracellular proteins expressed during the mineralization process of bone have high affinity for the hydroxyapatite mineral that makes up the majority of the crystalline component of bone mineral, including osteocalcin (9, 10) and intrinsically-disordered proteins (IDPs) such as osteonectin (8), osteopontin (25) and dentin matrix protein-1 (DMP1) (26), although DMP1 has been found to require the presence of another PILP to form intrafibrillar mineral (6). Osteocalcin is the most abundant of these calcium binding osteogenic proteins and has been found associated with both the exterior and interior of collagen fibrils in calcifying turkey tendon (27), leading osteocalcin to be considered a candidate biomolecule for the delivery of calcium ions into collagen fibrils. A biomolecule delivering sufficient calcium ions to make one 2×25×40 nm hydroxyapatite crystal must deliver a minimum of 1.6×10^5^ Ca^2+^ ions. Osteocalcin has five Ca^2+^ binding sites, so 3.2×10^4^ molecules of osteocalcin are required to build a single mineral crystal. Each monomer unit in PAR (molecular mass 0.6 kDa) has a theoretical Ca^2+^ capacity of up to four Ca^2+^ ions, so on the order of 2000 PAR molecules of the same molecular mass as one osteocalcin molecule (11 kDa, 20 ADP-ribose monomers per PAR polymer) are required to build a similar mineral crystal. Thus approximately 16 times less biomass is needed to deliver the necessary calcium via PAR than via osteocalcin. The other osteogenic proteins have similar calcium binding capacities to osteocalcin, e.g. osteonectin (SPARC) has a maximum of one Ca^2+^ per 3.8 kDa. PAR thus represents a significantly more energy-efficient route for osteoblasts to deliver calcium to collagen fibrils than osteocalcin.

Mice in which the two genes encoding for osteocalcin (*Bglap* and *Bglap2*) are knocked out show no deficiency in bone mineral density, but rather dysfunctional organization of the hydroxyapatite crystalline platelets with respect to the collagen fibril axis (9). This suggests that osteocalcin is not involved in delivery of calcium ions to collagen fibrils but may be involved in building the 3D architecture of bone mineral. We speculate that osteocalcin and osteogenic IDPs may bind to PAR-induced intrafibrillar mineral crystals to form the architecture of the mineral crystals that extend between collagen fibrils.

The PARP enzyme responsible for extracellular PAR has yet to be elucidated. PARylation of the osteogenic master transcription factor, RUNX2 has been linked to osteogenic differentiation of VSMCs under stress conditions (28) whilst PARP2 was found to be upregulated in calcification conditions in vitro in both osteoblasts and VSMCs (12), suggesting that both the major PAR-synthesis enzymes are important in calcification. PARP1 has been reported as primarily confined to cell nuclei (29) and to play roles in regulation of transcription (30) whilst PARP2 is found in both cell nuclei and the cytoplasm, and its roles in the cytoplasm have yet to be elucidated. Clearly it will be important to determine which PARP enzyme is responsible for synthesis of the extracellular PAR in bone and vascular calcification to define drug targets in bone and vascular calcification pathologies; currently, PARP2 would appear to be the strongest candidate, having a known presence in the cytoplasm.

## Conclusion

Our data suggest that both collagen type I C-terminal telopeptides and periodically ordered collagen fibrils may be necessary for functional collagen calcification in bone. This raises the questions of what happens if the C-terminal telopeptides are mutated or chemically altered or the ordered molecular organization in collagen fibrils is disrupted. Collagen Tyr residues have been found to be oxidized in ex vivo tissues (31). The only Tyr residues in collagen type I in most vertebrates are in the N- and C-terminal telopeptides, implying that such “accidental” chemistry could be expected to affect the C-terminal PAR binding sites and impact initiation of collagen calcification. We have previously shown that the molecular organization in collagen fibrils is disrupted by non-enzymatic glycation, a ubiquitous feature of ageing collagen (32). It is possible that pathologies involving dysfunctional bone mineral may stem from these types of mechanism, if so new drug targets for these pathologies may be revealed.

## Materials and Methods

### Transmission Electron Microscopy

All images were obtained using a Thermo Scientific (FEI) Talos F200X G2 field emission gun TEM operating at 200 keV.

OptiCol™ Rat Collagen Type I (3.9 mg/mL in 0.02M acetic acid) was used in all experiments. The supplied neutralizing solution was added in a 1:9 ratio to the required volume of collagen/acetic acid solution, and the neutralized mixture was left to form fibrils at 4°C overnight.

0.4 mg of pepsin (Promega) was added to 1 mL collagen/acetic acid, mixed, and left at 4°C. Aliquots were taken out after 48 hours, to which 1:9 neutralizing solution was added. The neutralized mixtures were left at 4°C to fibrillize.

Custom made, EDTA free (PAR) was obtained from BioTechne/Trevigen and had a concentration of 100 mM (56 mg/mL in 10 mM TRIS buffer, pH 8.0). Aliquots of PAR solution were added to an equal volume of a 1 mM CaCl_2_ solution prepared in water to give a 0.5 mM CaCl_2_ solution with PAR. PAR-Ca solutions were left at room temperature for 20-30 minutes prior to imaging.

For each sample, 2.3 μL of collagen suspension (raw or pepsin digested) was added to a TEM grid for 90 seconds, washed with water then partially blotted. 2.3 μL PAR-Ca solution was added to the same grid for 60 seconds, which was then stained with 2% uranyl acetate solution for 40 seconds, then blotted and imaged.

To analyze binding specificity of PAR-Ca droplets to collagen, the sub-band of collagen at the center of each PAR-Ca droplet was counted as the binding location for that droplet. A χ^2^ test was performed on the distribution of sub-band bindings between the two data sets.

### Sequence Conservation

Sequences were obtained from the National Centre for Biotechnology Information (NCBI) Reference Sequence Database for collagen I α1 and α2 chains of vertebrates. α1a sequences for Actinopterygii were combined with the available α1 sequences for all other vertebrate classes. Sequences were manually curated to remove outliers in terms of sequence length and other incorrectly assigned sequences, and isoforms were removed so that each species had at most one sequence. A total of 392 and 454 sequences remained, for α1 (including α1a) and α2 respectively. These were aligned using COBALT (COnstraint-Based multiple ALignment Tool) (33), and conservation at each site was measured.

Sequence logos were generated using WebLogo (34).

### Solution-State NMR

All NMR data was acquired on Bruker AVIII 500 MHz spectrometers equipped with either a BBO Smartprobe or DCH Cryoprobe. Spectra were obtained at 278K unless otherwise specified.

Custom model peptides were obtained from Bioserv UK (Ac-SAGFDFSFLPQPPQEKAHDGGRYYRA (α1) and Ac-GVSGGGYDFGYDGDFYRA (α2)). Adenosine diphosphate ribose (ADP-ribose) was obtained from Merck as a sodium salt.

Peptide solutions contained 1mM peptide, 0.71 mg/mL sodium trimethylsilylpropanesulfonate (DSS) as a ^1^H chemical shift reference, 10 mM ammonium bicarbonate buffer and 1/7 D_2_O by volume.

For assignment, a DCH Cryoprobe was used. 1D ^1^H, COSY, TOCSY, NOESY and ROESY spectra were obtained at 278K.

For titration with ADP-ribose, 1D ^1^H spectra were obtained using a BBO Smartprobe, with water suppression achieved using excitation sculpting with gradient pulses.(35) After each acquisition, 2-4 μL of neutral 350 mM ADP-ribose solutions in 10 mM aqueous sodium hydroxide was added to the NMR tube which was then shaken. Relative concentration of ADP-ribose to peptide was calculated by integrating the adenosine ribose 1’ proton signal between 6.1-6.2 ppm and comparing to the 6H leucine (α1) and 6H valine (α2) signals at around 0.9 ppm in each peptide. Titrations were recorded at 278K, 288K, and 298K.

## Acknowledgements

Funded by the European Union (ERC, EXTREME 101019499). Views and opinions expressed are however those of the author(s) only and do not necessarily reflect those of the European Union or the European Research Council Executive Agency. Neither the European Union nor the granting authority can be held responsible for them.

EPSRC Underpinning Multi-User Equipment Call (EP/P030467/1) for funding the TEM.

## Notes

### Competing Interest Statement

The authors have declared no competing interest.

