## Supplementary material for "Poly(ADP-ribose) binding sites on collagen I fibrils for nucleating intrafibrillar bone mineral": SI

| Proton environment | NH | α | β | γ | δ | ε |
|---|---|---|---|---|---|---|
| α1 R22 | 8.07 | 4.20 | 1.63 | 1.38 | 3.09 | |
| α1 Y23 | 8.30 | 4.55 | 2.98, 2.85 | | 7.03 | 6.76 |
| α1 Y24 | 8.13 | 4.46 | 2.91 | | 7.04 | 6.77 |
| α1 R25 | 8.29 | 4.20 | 1.77,1.65 | 1.55 | 3.15 | |
| α2 Y7 | 8.15 | 4.44 | 2.86, 2.77 | | 6.91 | 6.74 |
| α2 Y11 | 8.08 | 4.51 | 3.01, 2.94 | | 7.05 | 6.75 |
| α2 Y16 | 8.18 | 4.39 | 2.92 | | 7.06 | 6.80 |
| α2 R17 | 7.93 | 4.20 | 1.75, 1.64 | 1.54 | 3.13 | |

*Table S1: $^{1}H$ chemical shifts of the model peptides' amino acids investigated in this work at 278K in 6:1 $H_2O$:$D_2O$ referenced to sodium trimethylsilylpropanesulfonate (DSS)*

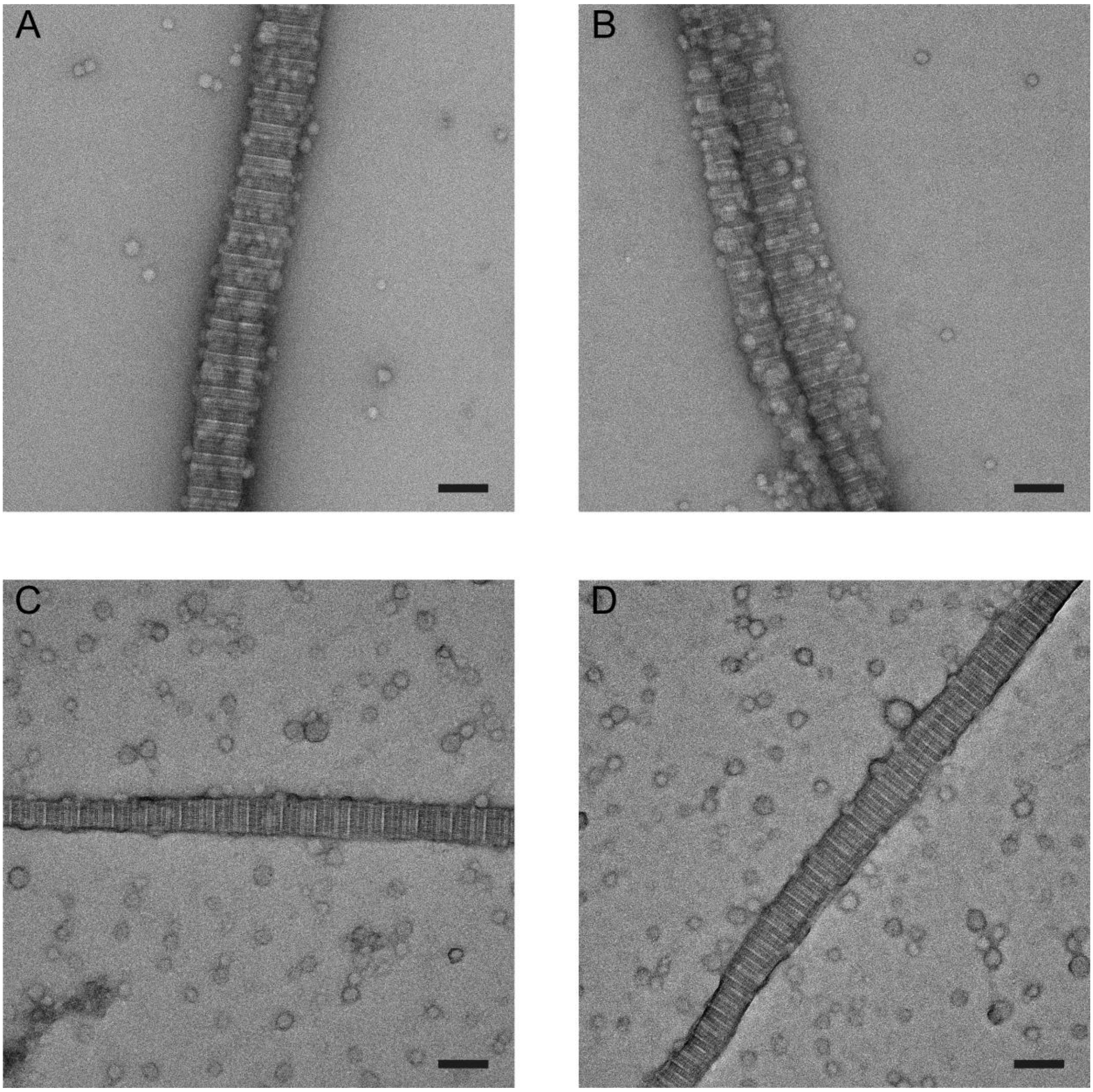


*Figure S1: TEM images of PAR-Ca droplets bound to an untreated collagen fibril (A, B) and a collagen fibril formed from collagen that had been degraded with pepsin for 48 hours prior to fibrillization (C, D), showing decreased binding affinity of PAR-Ca droplets to pepsin-digested collagen fibrils (scale bars 100 nm).*

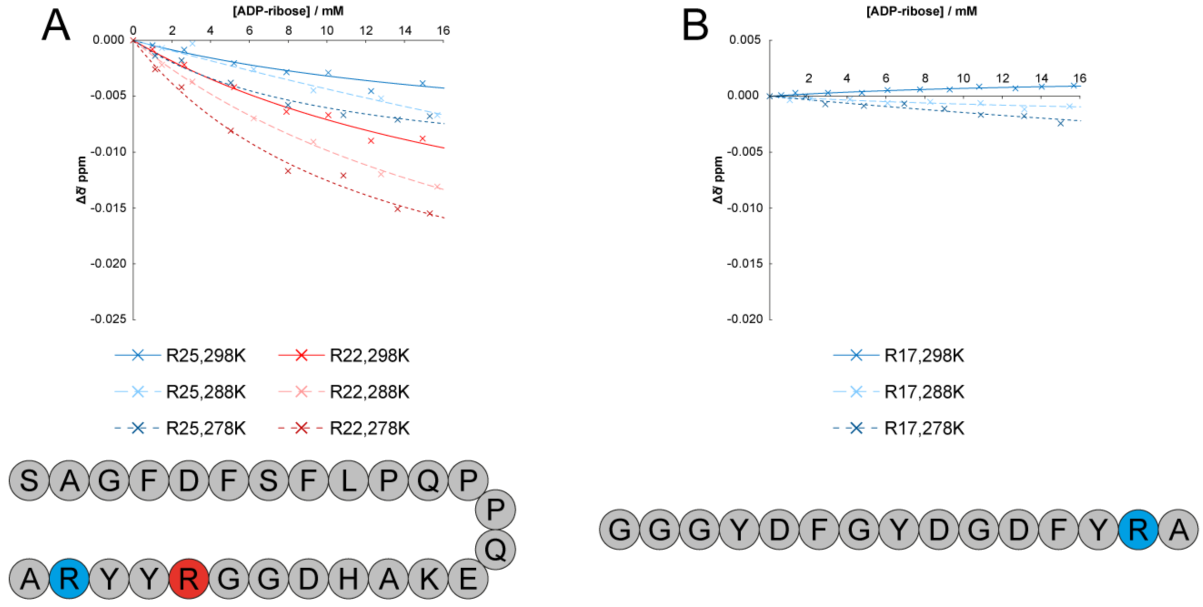


*Figure S2: Plots showing the change in chemical shift of arginine δ protons in the α1 (A) and α2 (B) model peptides upon addition of ADP-ribose. Approximate binding curves have been fitted according to equation {1}.*

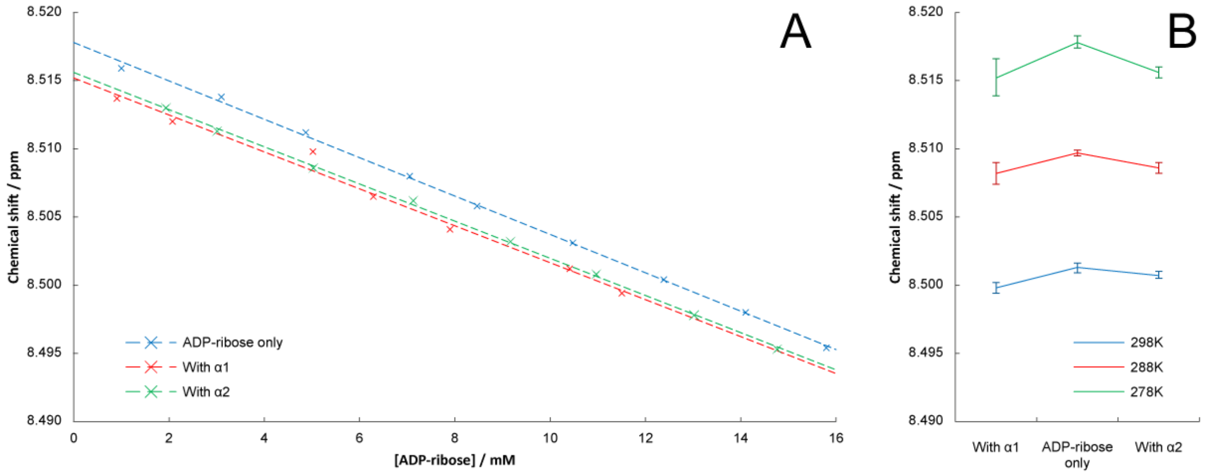


*Figure S3: A: Plot of ADP-ribose H8 chemical shift changes at 278K as the concentration of ADP-ribose is increased, in the presence or absence of either of the two model peptides. Straight lines of best fit plotted and extrapolated back to zero ADP-ribose concentration, which is the theoretical chemical shift of the adenine H8 proton of an infinitely dilute solution of ADP-ribose such that there is no adenine-adenine π-stacking. B: Plot showing that the extrapolated zero ADP-ribose chemical shift of adenine H8 is decreased in the presence of each of the model peptides.*
